# CUA: a Flexible and Comprehensive Codon Usage Analyzer

**DOI:** 10.1101/022814

**Authors:** Zhenguo Zhang

**Author notes:** Correspondence: University of Rochester, River Campus Box 270211 Rochester, New York 14627-0211; or.

## Abstract

Codon usage bias (CUB) is pervasive in genomes. Studying its patterns and causes is fundamental for understanding genome evolution. Rapidly emerging large-scale RNA and DNA sequences make studying CUB in many species feasible. Existing software however is limited in incorporating the new data resources. Therefore, I release the software CUA which can compute all popular CUB metrics, including *CAI, tAI, Fop, ENC*. More importantly, CUA allows users to incorporate user-specific data, such as tRNA abundance and highly expressed genes from considered tissues; this flexibility enables computing CUB metrics for any species with improved accuracy. In sum, CUA eases codon usage studies and establishes a platform for incorporating new metrics in future. CUA is available at http://search.cpan.org/dist/Bio-CUA/ with help documentation and tutorial.

## Introduction

One amino acid can be encoded by more than one synonymous codon, and synonymous codons are unevenly used. In particular, some codons are used more often than their synonymous ones in highly expressed genes (Sharp and Li, 1987). To measure the unevenness of codon usage, multiple metrics of codon usage bias (CUB) have been developed, such as Codon Adaptation Index (*CAI*) (Sharp and Li, 1987), tRNA Adaptation Index (*tAI*) (dos Reis, et al., 2004), Frequency of optimal codons (*Fop*) (Ikemura, 1981), and Effective Number of Codons (*ENC*) (Wright, 1990). With assumptions, all of these metrics except *ENC* also measure translation efficiency: the larger the metrics, the more efficiently translated the codons are.

CUB can be calculated at two levels: codon and sequence. At the codon level, the metrics measure the relative translation efficiency among codons. For example, codons preferred in highly expressed genes (Sharp and Li, 1987) or matching more cognate tRNAs (Ikemura, 1981) are thought to be translated more efficiently than their synonymous alternatives. At the sequence level, the metrics measure the overall translation efficiency (or, for *ENC*, codon usage unevenness only) of a sequence, and for *CAI* and *tAI*, they are computed by averaging (usually taking the geometric mean) the codon-level values of the sequence’s constituent codons (dos Reis, et al., 2004; Sharp and Li, 1987). Note that *ENC* can be computed only at the sequence level.

CUB is observed in prokaryotes and eukaryotes (Chen, et al., 2004; Vicario, et al., 2007), and its patterns and causes are informative about translation regulation and natural selection (Akashi, 1994; Bentele, et al., 2013; Bulmer, 1991; Eyre-Walker, 1991; Goodman, et al., 2013; Hambuch and Parsch, 2005; Plotkin and Kudla, 2011; Pop, et al., 2014; Qian, et al., 2012; Singh, et al., 2005). Previously, CUB was studied in a few species where genomes or gene expression data were available. The growing huge amount of DNA and RNA sequence data make studying CUB in many species feasible. For example, *tAI* can be computed for any sequenced genomes using tRNA copy numbers to approximate tRNA abundance (dos Reis, et al., 2004). For another example, more accurate *CAI* can be computed by using the highly expressed genes from the considered tissues as the reference gene set other than using a general gene set (Qian, et al., 2012).

Existing software packages, however, are limited in incorporating user-specific data and thus usable only for certain circumstances. codonW (http://codonw.sourceforge.net/) is great for computing sequence-level *ENC*, *CAI*, and *Fop* for a few species whose codon-level CUB values have been integrated, but inconvenient for other species whose codon-level values are not implemented. codonR (http://people.cryst.bbk.ac.uk/∼fdosr01/tAI/) can compute tAI for *Escherichia coli*, but for other species one need specify tRNA abundance in a not well-documented format, causing inconvenience. DAMBE (Xia, 2013) is flexible in calculating CAI by accepting user-specific codon tables, but this flexibility can be further improved (see below).

To advance codon usage studies, I have developed a new software package CUA, short for Codon Usage Analyzer. The package implements the four popular CUB metrics and is flexible in incorporating user-specific data.

## Implementation and usage

The package provides both PERL modules and ready-to-use programs, and can be easily integrated into large-scale data analysis pipeline. Users can write new programs based on the modules or use the provided programs to accomplish their work. The package should work in any operation system with PERL installed. The package is deposited into Comprehensive Perl Archive Network (CPAN) at http://search.cpan.org/dist/Bio-CUA/, and can be installed by typing ‘cpan Bio::CUA’ in the command line.

Overall, the package comprises three main modules: Bio::CUA::CUB::Builder, Bio::CUA::CUB::Calculator and Bio::CUA::Summarizer. The former two modules compute CUB metrics at the codon and sequence levels, respectively, while Bio::CUA::Summarizer supports the computation by providing common functions of sequence processing. See http://search.cpan.org/dist/Bio-CUA/ for more modules included in the package.

One main advantage of CUA is its flexibility in setting user-specific parameters. For example, it can use tRNA abundance of any species in a simple format for computing *tAI*. It also accepts both codon tables and sequences for *CAI* calculation, and allows specifying user-specific highly expressed genes other than using a general reference set. It is also flexible in specifying genetic code tables. In the following paragraphs, I will demonstrate some of its flexibilities by giving an example of calculating CUB metrics for genes in the fruitfly *Drosophila melanogaster*. The commands and data are listed in supplementary Text S1.

First, I compute the codon-level *tAI* and *CAI* using the programs *tai_codon.pl* and *cai_codon.pl*, respectively. For *tAI*, I download tRNA copy numbers from GtRNAdb (http://gtrnadb.ucsc.edu/) (Chan and Lowe, 2009) for *D. melanogaster*, sum them up for each anticodon (tRNA abundance can be used instead if available), and use it as input to the program *tai_codon.pl* for computing the codon-level *tAI*. For *CAI*, I extract the top 200 highly expressed genes as the reference gene set (as in Qian, et al., 2012) based on mRNA expression levels from the *Drosophila* S2 cell line (Zhang, et al., 2010), download protein-coding sequences from FlyBase (St Pierre, et al., 2014), and use these two datasets as input to the program *cai_codon.pl* for computing the codon-level *CAI*.

Second, I compute the sequence-level CUB metrics using the program *cub_seq.pl*. I feed the program with the codon-level *tAI* and *CAI* calculated above and also specify the *ENC* option, so that the sequence-level *tAI*, *CAI* and *ENC* are calculated along with other parameters such as counts of amino acids and GC content. Users can also refer to the tutorial at http://search.cpan.org/dist/Bio-CUA/lib/Bio/CUA/Tutorial.pod for computing other CUB metrics.

## CAI and ENC variants

In addition to the original *CAI* (Sharp and Li, 1987), CUA also implements two CAI variants, *mCAI* and *bCAI*. They differ from the original one in normalizing relative synonymous codon usage (RSCU) (Sharp and Li, 1987). *mCAI* and *bCAI* take into account RSCUs expected from even codon usage and from the background data (*e.g.*, RSCUs of lowly expressed genes), respectively. See Text S2 for details. I compare these *CAI* variants by correlating them with mRNA expression and translation efficiency (determined by ribosome profiling technique (Ingolia, et al., 2009)) using the data of the S2 cell line (Dunn, et al., 2013; Zhang, et al., 2010). The better metric is expected to correlate more strongly. As shown in Table 1, *bCAI* correlates with mRNA expression and translation efficiency more strongly than CAI and *mCAI* do, and the latter two perform somewhat equivalently. Therefore, *bCAI* performs better when predicting mRNA expression levels or translation efficiency, although only slightly because most comparisons are not statistically significant.

**Table 1.** Correlation of CAI metric variants with mRNA expression and translation efficiency in *Drosophila* S2 cells.

|  |  | *CAI* | *mCAI* | *bCAI* |
| --- | --- | --- | --- | --- |
| Correlation with mRNA expression | Spearman's Rho | 0.402713 | 0.390343 | 0.421782 |
|  | Spearman's *P* | < 2.2e-16 | < 2.2e-16 | < 2.2e-16 |
|  | Pearson's R | 0.473567 | 0.471513 | 0.504091* |
|  | Pearson's *P* | < 2.2e-16 | < 2.2e-16 | < 2.2e-16 |
| Correlation with translation efficiency | Spearman's Rho | 0.090826 | 0.070669 | 0.103919 |
|  | Spearman's *P* | 6.50E-11 | 3.81E-07 | 7.53E-14 |
|  | Pearson's R | 0.069513 | 0.056945 | 0.080018 |
|  | Pearson's *P* | 5.89E-07 | 4.31E-05 | 8.83E-09 |

In order to reduce sampling errors in RNA sequencing, genes with mRNA RPKM > 10 are used (5153 genes in total). Using RPKM > 1 does not alter the conclusion. \* indicates the correlation coefficients significantly differ between *bCAI* and *CAI* ( $P < 0.05$ ). Translational efficiency is measured by ribosome profiling technology and obtained from the literature (Dunn, et al., 2013).

For *ENC*, CUA implements both the original (Wright, 1990) and nucleotide-composition-corrected *ENC* (Novembre, 2002), and tries to estimate missing *F* values using a different way (see Text S2 for details). I compare these *ENC* variants using the protein coding genes in *D. melanogaster* and using the nucleotide compositions from each gene’s introns to correct nucleotide compositions. The results show that the method for estimating missing *F* values has little effect on *ENC* values because most sequences are long enough to obtain all *F* values, and that, instead, correcting nucleotide composition has strong effect on *ENC* values as the corrected and non-corrected *ENC*s are moderately correlated (Fig. S3). Surprisingly, we find the corrected ENCs generally perform worse than non-corrected ones in terms of correlation with mRNA expression levels and translation efficiency (Table 2), though ENC is designed only to measure codon usage unevenness other than to predict mRNA expression or translation efficiency.

**Table 2.** Correlation of ENC variants with mRNA expression, translation efficiency, and *tAI* using data from *Drosophila* S2 cells.

|  |  | *ENC*<sup>a</sup> | *ENC_r*<sup>a</sup> | *ENCp*<sup>a</sup> | *ENCp_r*<sup>a</sup> |
| --- | --- | --- | --- | --- | --- |
| Correlation with mRNA expression | Spearman's Rho | -0.34332 | -0.3452 | -0.24139 | -0.23952 |
|  | Spearman's *P* | < 2.2e-16 | < 2.2e-16 | < 2.2e-16 | < 2.2e-16 |
|  | Pearson's R | -0.41732 | -0.42122 | -0.26769 | -0.27255 |
|  | Pearson's *P* | < 2.2e-16 | < 2.2e-16 | < 2.2e-16 | < 2.2e-16 |
| Correlation with translation efficiency | Spearman's Rho | -0.04274 | -0.04545 | -0.04954 | -0.05164 |
|  | Spearman's *P* | 0.002151 | 0.001101 | 0.008827 | 0.006336 |
|  | Pearson's R | -0.05374 | -0.05379 | 0.010019 | 0.010623 |
|  | Pearson's *P* | 0.000114 | 0.000112 | 0.5966 | 0.5747 |
| Correlation with *tAI* | Spearman's Rho | -0.69335 | -0.69023 | -0.54936 | -0.54894 |
|  | Spearman's *P* | < 2.2e-16 | < 2.2e-16 | < 2.2e-16 | < 2.2e-16 |
|  | Pearson's R | -0.69955 | -0.69525 | -0.56535 | -0.56504 |
|  | Pearson's *P* | < 2.2e-16 | < 2.2e-16 | < 2.2e-16 | < 2.2e-16 |

a: *ENCp* and *ENC* stand for the metrics with and without nucleotide composition corrected, respectively; and *ENCp\_r* and *ENC\_r* are the corresponding versions with missing *F* values estimated using Equation 4 in Text S2. In order to reduce sampling errors in RNA sequencing, genes with mRNA RPKM > 10 are used (5153 genes in total). Using RPKM > 1 does not alter the conclusion.

## Conclusions and Future Directions

In sum, CUA is flexible in incorporating users’ data and able to compute all popular CUB metrics for any species, which will facilitate CUB studies.

Additionally, I plan to implement more CUB metrics and add a graphic user interface based on users’ feedback. I also expect expanding the package by more contributors because of the great collaborating environment through CPAN.

## Supplementary information

Supplementary Material is available online.

## Acknowledgements

I am very grateful to Dr. Daven Presgraves for supporting this project and also thank Dr. Harry Stern and other Bluehive staffs at University of Rochester for their excellent support for computation.

## References

Akashi, H. (1994) Synonymous codon usage in Drosophila melanogaster: natural selection and translational accuracy. Genetics, 136(3), 927–935.

Bentele, K., et al. (2013) Efficient translation initiation dictates codon usage at gene start. Mol Syst Biol, 9, 675.

Bulmer, M. (1991) The selection-mutation-drift theory of synonymous codon usage. Genetics, 129(3), 897–907.

Chan, P.P. and Lowe, T.M. (2009) GtRNAdb: a database of transfer RNA genes detected in genomic sequence. Nucleic Acids Res, 37(Database issue), D93–97.

Chen, S.L., et al. (2004) Codon usage between genomes is constrained by genome-wide mutational processes. Proc Natl Acad Sci U S A, 101(10), 3480–3485.

dos Reis, M., Savva, R. and Wernisch, L. (2004) Solving the riddle of codon usage preferences: a test for translational selection. Nucleic Acids Res, 32(17), 5036-5044.

Dunn, J.G., et al. (2013) Ribosome profiling reveals pervasive and regulated stop codon readthrough in Drosophila melanogaster. Elife, 2, e01179.

Eyre-Walker, A.C. (1991) An analysis of codon usage in mammals: selection or mutation bias? J Mol Evol,33(5), 442–449.

Goodman, D.B., Church, G.M. and Kosuri, S. (2013) Causes and effects of N-terminal codon bias in bacterial genes. Science, 342(6157), 475–479.

Hambuch, T.M. and Parsch, J. (2005) Patterns of synonymous codon usage in Drosophila melanogaster genes with sex-biased expression. Genetics, 170(4), 1691–1700.

Ikemura, T. (1981) Correlation between the abundance of Escherichia coli transfer RNAs and the occurrence of the respective codons in its protein genes. J Mol Biol,146(1), 1–21.

Ingolia, N.T., et al. (2009) Genome-wide analysis in vivo of translation with nucleotide resolution using ribosome profiling. Science, 324(5924), 218–223.

Novembre, J.A. (2002) Accounting for background nucleotide composition when measuring codon usage bias. Mol Biol Evol,19(8), 1390–1394.

Plotkin, J.B. and Kudla, G. (2011) Synonymous but not the same: the causes and consequences of codon bias. Nat Rev Genet,12(1), 32–42.

Pop, C., et al. (2014) Causal signals between codon bias, mRNA structure, and the efficiency of translation and elongation. Mol Syst Biol, 10, 770.

Qian, W., et al. (2012) Balanced codon usage optimizes eukaryotic translational efficiency. PLoS Genet, 8(3), e1002603.

Sharp, P.M. and Li, W.H. (1987) The codon Adaptation Index-a measure of directional synonymous codon usage bias, and its potential applications. Nucleic Acids Res,15(3), 1281–1295.

Singh, N.D., Davis, J.C. and Petrov, D.A. (2005) X-linked genes evolve higher codon bias in Drosophila and Caenorhabditis. Genetics, 171(1), 145–155.

St Pierre, S.E., et al. (2014) FlyBase 102-advanced approaches to interrogating FlyBase. Nucleic Acids Res, 42(Database issue), D780–788.

Vicario, S., Moriyama, E.N. and Powell, J.R. (2007) Codon usage in twelve species of Drosophila. BMC Evol Biol, 7, 226.

Wright, F. (1990) The ‘effective number of codons’ used in a gene. Gene, 87(1), 23–29.

Xia, X. (2013) DAMBE5: a comprehensive software package for data analysis in molecular biology and evolution. Mol Biol Evol, 30(7), 1720–1728.

Zhang, Y., et al. (2010) Expression in aneuploid Drosophila S2 cells. PLoS Biol, 8(2), e1000320.

